# The Effect of Selected Coffee Constituents on Mitochon-Drial Dysfunction In An Early Alzheimer Disease Cell Model

**DOI:** 10.1101/2023.04.20.537643

**Authors:** Lukas Babylon, Micha T. Limbeck, Gunter P. Eckert

## Abstract

Alzheimer disease (AD) is an emerging medical problem worldwide without any cure yet. By 2050, more than 152 million people will be affected. AD is characterized by mitochondrial dys-function (MD) and increased amyloid beta (Aβ) levels. Coffee is one of the most commonly consumed beverages. It has many bioactive and neuroprotective ingredients of which caffeine (Cof), kahweohl (KW) and cafestol (CF) shows a variety of pharmacological properties such as anti-inflammatory and neuroprotective effects. Effects of Cof, KW, and CF were tested in a cel-lular model of AD on MD and Aβ. SH-SY5Y-APP695 cells were incubated with 50µM Cof, 1µM CF and 1µM KW for 24h. The energetic metabolite ATP was determined using a luciferase-catalyzed bioluminescence assay. The activity of mitochondrial respiration chain complexes was assessed by high-resolution respirometry using a Clarke electrode. Expression levels genes were deter-mined using quantitative real-time polymerase chain reaction (qRT-PCR). The levels of amyloid β-protein (Aβ_1-40_) were measured using homogeneous time-resolved fluorescence (HTRF). ROS levels, cAMP levels, and peroxidase activity were determined using a fluorescence assay. The combination of Cof, KW and CF significantly increased ATP levels. The combination had neither a significant effect on MMP, on activity of respiration chain complexes, nor on Aβ_1-40_ levels. cAMP levels were slightly increased after incubation with the combination, but not the peroxi-dase activity. Pyruvate levels and the lactate-pyruvate-ration but not lactate levels were significantly enhanced. No effect was seen on the expression level of lactate dehydrogenase and py-ruvate dehydrogenase kinase. In some experiments we have tested the single substances. They showed significant results especially in ATP, lactate and pyruvate values compared to the con-trol. The combinations have a lesser effect on mitochondrial dysfunction in cells and none on Aβ production. Whereas ATP levels and pyruvate levels were significantly increased. This suggests a change in glycolysis in neuronal cells harbouring human genes relevant for AD.

## 1. Introduction

Alzheimer’s disease (AD) is a neurodegenerative disease belonging to the class of dementias. It manifests itself through a progressive loss of function of neurons of the central nervous system [1]. Dementia affects 5-7% of people over 60 in developed countries [2], of which Alzheimer’s disease, discovered by German physician Alois Alzheimer in 1906, is the most common manifestation, accounting for 60% [1]. Clinical symptoms include memory loss as well as language and orientation problems [2]. One possible cause of AD is a manifestation of mitochondrial dysfunction (MD) as an early event of AD [3,4]. Almost all mitochondrial functions are impaired in AD [5,6]. First signs of MD are a reduction in key enzymes of in the oxidative metabolism [7,8] and glucose consumption [9]. The limited function of the electron transport chain (ETC) is the reason for the decrease activity of complexes IV and I. These complications lead to a lower membrane potential (MMP) and to reduced ATP levels [10,11]. A further possible trigger of the emergence of AD is the overexpression of amyloid-beta (Aβ) [12]. An imbalance between production and removal leads to accumulation and aggregation in the brain, this leads to inflammatory responses, production of reactive oxygen species (ROS) and loss of neurons, this leads to MD, dementia and Alzheimer disease [13]. An important antioxidant enzyme involved in AD is peroxidase, which catalyzes the oxidation of organic and non-organic substances using H2O2. In this process, it is supposed to protect against the harmful effects of physiological accumulation of ROS. A change in peroxidase activity could induce effects on ROS levels. Another impairment in AD is cyclic adenosine monophosphate (cAMP), which is one of the nucleotides and serves as a second messenger in the body for signaling metabolic pathways and hormone effects. [14,15]. The signaling cascade consisting of cAMP/PKA/CREB is considered to be generally important for the processes and functioning in learning and memory [16]. Performance deficits may be mediated by deficits in the signaling cascade. One trigger of this impairment could be Aβ toxicity [17]. Up to now, there is no cure for it, which is why prevention and therapy are in the focus of attention. A possibility for this offer might be coffee.

Today, there is a high consumption of coffee worldwide, which in Germany is 150 l per capita per year [18]. Due to a high diversity of bioactive ingredients, the beverage has been the subject of research for many years regarding its influence on human health. Caffeine has been the focus of attention due to its stimulant effects. In addition, many other secondary plant compounds with bioactive effects are now known in coffee. These potentially health-promoting effects are based on cardioprotective, hepatoprotective and neuroprotective properties [19]. Roasted coffee contains a variety of bioactive ingredients. Besides caffeine, coffee contains other diterpenes in the form of fatty acid esters, with cafestol and kahweol making up the largest proportion [20].

Depending on the coffee variety, the composition of cafestol and kahweol differs. Arabica coffee contains cafestol and kahweol, whereas robusta contains cafestol and small amounts of kahweol [21]. The effects of caffeine are due to mechanisms of action on nerve cells in the brain, where the pathological changes during AD also occur [22]. Coffee and caffeine intake is associated with a lower risk of developing dementia and AD through a wide variety of mechanisms [23–25]. The data situation on kahweol and cafestol is limited, but there are some evidence that both substances also have a neuroprotective effect [26–28]. In this study, we focused on the combination of caffeine, kahweol and cafestol in small doses on an early AD cell model.

1. 2. Materials and Methods

### 2.1 Cell culture

Human neuroblastoma SH-SY5Y cells were cultured at 37 °C under an atmosphere of 5% CO_2_ in DMEM supplemented with 10% (v/v) heat-inactivated fetal calf serum, 60µg/mL streptomycin, 60units/mL penicillin, 0.3mg/mL hygromycin, MEM non-essential amino acids, and 1mM sodium pyruvate 1%. SH-SY5Y cells were stably transfected with DNA constructs harboring human wild-type APP_695_ (APP_695_) (for details; please refer to [29]). Cells were passaged every 3 days and were used for experiments when they reached 70-80% confluence.

### 2.2 Cell treatment

Cells were incubated with 1µM kahweol (KW), 1µM cafestol (Caf) and 50µM caffeine (Cof) (KCC) for 24h. The solution medium DMSO served as control.

### 2.3 ATP Measurement

A bioluminescence assay was used to determine the ATP levels, which is based on the production of light from ATP and luciferin in the presence of luciferase. The test was performed using the ATPlite Luminescence Assay System (PerkinElmer, Rodgau, Germany) according to the previously published protocol [30].

### 2.4 MMP Measurement

Mitochondrial membrane potential (MMP) was measured using the fluorescence dye rhodamine-123 (R123). Cells were incubated for 15 min with 0.4 µM R123 and centrifuged at 750 x g for 5 min before being washed with Hank’s Balanced Salt Solution (HBSS) buffer supplemented with Mg^2+^, Ca^2+,^ and HEPES. The Cells were suspended with fresh HBSS before they were evaluated by measuring the R123 fluorescence. The excitation wavelength was set to 490 nm and the emission wavelength to 535 nm.

### 2.5 Cellular Respiration

Respiration in SH-SY5Y_695_ cells was assessed using an Oxygraph-2k (Oroboros, Innsbruck, Austria) and DatLab 7.0.0.2. The cells were treated according to a complex protocol developed by Dr Erich Gnaiger [31]. They were incubated with different substrates, inhibitors and uncouplers. First, cells were washed with PBS (containing potassium chloride 26.6 mM, potassium phosphate monobasic 14.705 mM, sodium chloride 1379.31 mM and sodium phosphate dibasic 80.59 mM) and scraped into mitochondrial respiration medium (MiRO5) developed by Oroboros [31]. Afterward, they were centrifuged, resuspended in MiRO5, and diluted to 10^6^ cells/mL. After 2mL of cell suspension was added to each chamber and endogenous respiration was stabilized, the cells were treated with digitonin (10µg/10^6^ cells) to permeabilize the membrane, leaving the outer and inner mitochondrial membrane intact. OXPHOS was measured by adding the complex I and II substrates malate (2mM), glutamate (10mM) and ADP (2mM), followed by succinate (10 mM). Gradual addition of carbonyl cyanide-4- before it is evaluated by measuring the R123 fluorescence (trifluoromethoxy) phenylhydrazone showed the maximum capacity of the electron transfer system. To measure complex II activity, rotenone (0.1mM), a complex I inhibitor, was injected. After that, oligomycin (2 µg/mL) was added to measure the leak respiration. Inhibition of complex III by the addition of antimycin A (2.5 µM) determined residual oxygen consumption, which was subtracted from all respiratory states. The activity of complex IV was measured by adding *N, N, N′, N′*-tetramethyl-*p*- phenylenediamine (0.5 mM) and ascorbate (2 mM). To measure the sodium autoxidation rate, azide (≥100mM) was added. Afterward, complex IV respiration was corrected for autoxidation. DMSO served as control.

### 2.6 Citrate Synthase Activity

Cell samples from respirometry measurements were frozen and stored at -80°C for the determination of citrate synthase activity. Samples were thawed while the reaction mix (0.1 mM 5,5′-dithiol-bis-(2-nitrobenzoic acid) (DTNB), 0.5 mM oxaloacetate, 50 μM EDTA, 0.31 mM acetyl coenzyme A, 5 mM triethanolamine hydrochloride and 0.1 M Tris-HCl) was mixed and heated at 30°C for 5 min. Afterward, 40 µl of samples were submitted in triplets and mixed with 110µl of the reaction mix. The absorption was measured at 412nm.

### 2.7 Aβ_1-40_ Determination

After 24 h incubation, the Aβ_1-40_ levels were determined in SH-SY5Y-APP_695_ cells using HTRF Amyloid-Beta 1-40 kit (Cisbio, Codolet, France). The protocol was the same as recently described [32]. Aβ concentrations were normalized against the protein content.

### 2.8. Peroxidase Activity

Peroxidase activation was measured using an AmplexTM Red Peroxidase Kit (Thermo Fisher Scientific, Waltham, MA, USA) according to the manufacturer’s instructions.

Cells were seeded in 96-well plates and incubated for 24 h.

### 2.9 Protein Quantification

Protein content was determined using a Pierce^TM^ Protein Assay Kit (Thermo Fisher Scientific, Waltham, MA, USA) according to the manufacturer’s instructions. Bovine serum albumin was used as a standard.

### 2.10. ROS Measurement

Cellular ROS production was determined using a DCFDA/H2DCFDA Cellular ROS Assay Kit (ab113851; Abcam, Cambridge, UK). Cells were incubated for 24h with our substances and then the manufacturer’s instructions were followed.

### 2.11 Pyruvate and Lactate Content

Frozen cells, which were previously harvested and incubated for 24h, were thawed at room temperature. Pyruvate and lactate concentrations were assessed using a pyruvate assay kit (MAK071, Sigma Aldrich, Darmstadt, Germany) and a lactate assay kit (MAK064, Sigma Aldrich, Darmstadt, Germany) according to the manufacturer’s instructions.

### 2.12 Real-Time qRT-PCR

Total RNA was isolated using the RNeasy Mini Kit (Qiagen, Hilden, Germany) according to the manufacturer’s guidelines. RNA was quantified using a Nanodrop^TM^ 2000c spectrometer (Thermo Fisher Scientific, Waltham, MA, USA). To remove residual genomic DNA, samples were treated with a TURBO DNA-free^TM^ kit according to the manufacturer’s instructions (Thermo Fisher Scientific, Waltham, MA, USA).

Complementary DNA was synthesized from 1 µg total RNA using an iScript cDNA Synthesis Kit (Bio-Rad, Munich, Germany). qRT-PCR was conducted using a CfX 96 Connect™ system (Bio-Rad, Munich, Germany). Primers were provided from Biomol (Hamburg, Germany). All primers are listed in Table 1. The cDNA aliquots were diluted 1:10 with RNase-free water (Qiagen, Hilden, Germany), and all samples were analyzed in triplicate. PCR cycling conditions were as follows: initial denaturation for 3 min at 95 °C, followed by 45 cycles at 95 °C for 10 s, 58 °C for 30 s (or 56 °C for 45 s, depending on the primer), and 72 °C for 29 s. Expression was analyzed with –(2ΔΔCq) using Bio-Rad CfX manager software. The normalization factor was calculated based on the geometric mean of the levels of multiple control genes of *ß-actin (ACTB), glyceraldehyde-3-phosphate dehydrogenase (GAPDH)*, and *phosphoglycerate kinase 1 (PGK1)* according to the MIQE guidelines [33]. No-template control served as an assay control to exclude impurities.

**Table 1.**
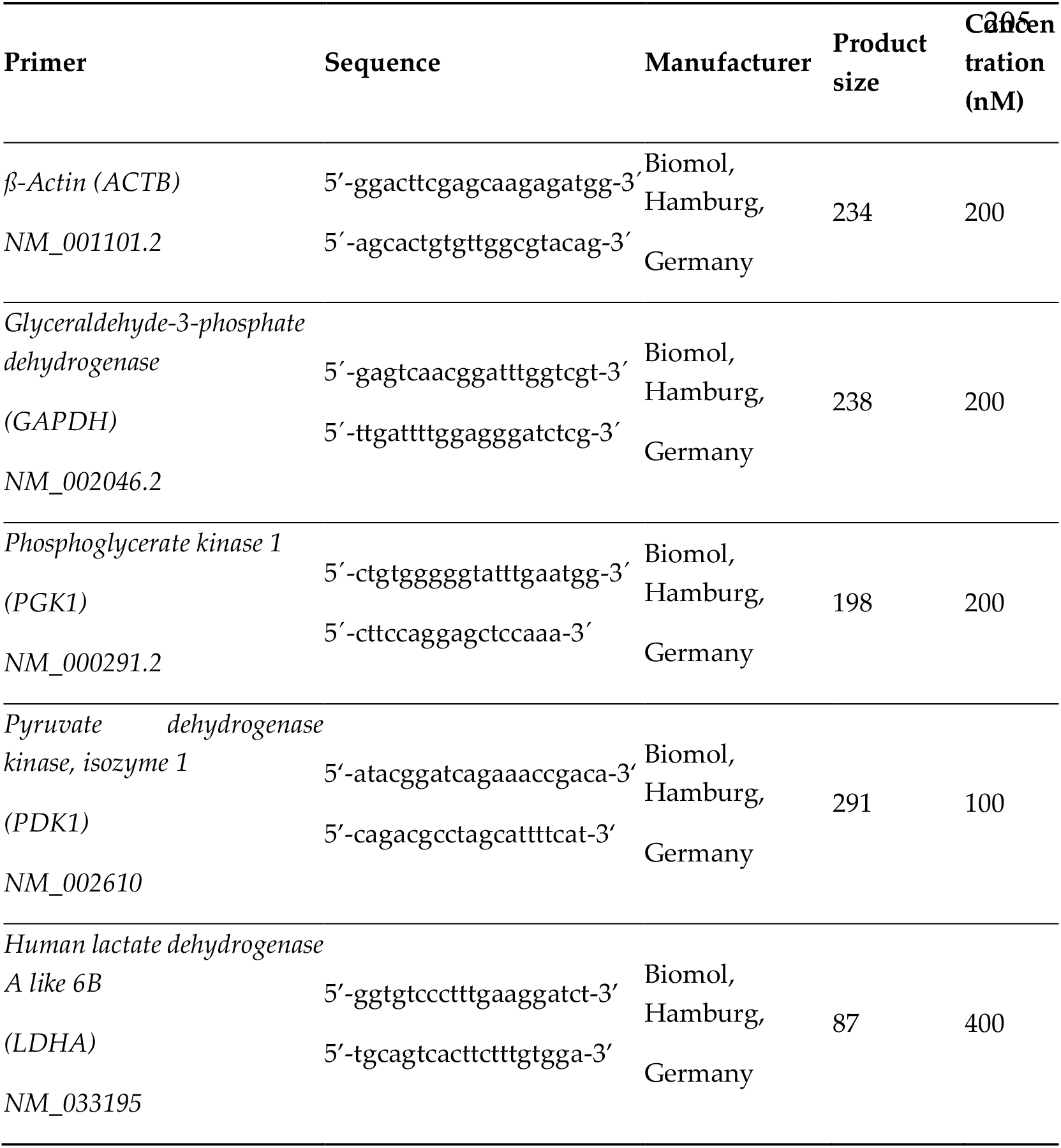
Oligonucleotide primer sequences, product sizes, and primer concentrations for quantitative real-time PCR.

### 2.13 cAMP level

Cellular peroxidase activity was determined using a cAMP Direct Immunoassays Kit (Abcam, Cambridge, UK). Cells were incubated for 24h with our substances and then the manufacturer’s instructions were followed.

### 2.14 Statistics

Unless otherwise stated, values are presented as mean ± standard error of the mean (SEM). Statistical analyses were performed by applying one-way ANOVA with Tukeýs multiple comparison post-hock test and student’s unpaired t-test (Prism 9.1 GraphPad Software, San Diego, CA, USA). Statistical significance was defined for *p* values * p < 0.05, ** p < 0.01, *** p < 0.001 and **** p < 0.0001.

## 3. Results

### 3.1 Mitochondrial functions

To find the right concentrations for our combination we tested different concentrations of Cof, CF and KW (SupFigure 1), as well as combinations of the three mentioned substances, for the best combination, for the effects on the ATP levels (SupFigure 2). The concentration of kahweol 1µM, cafestol 1µM and caffeine 50µM (KCC) contain all of our three substances and turned out to be the best. There the highest ATP values for a combination were achieved. We tested KCC on the mitochondrial functions, we incubated SH-SY5Y-APP695 cells for 24h with all three of the substances combined.

**Figure 1:**
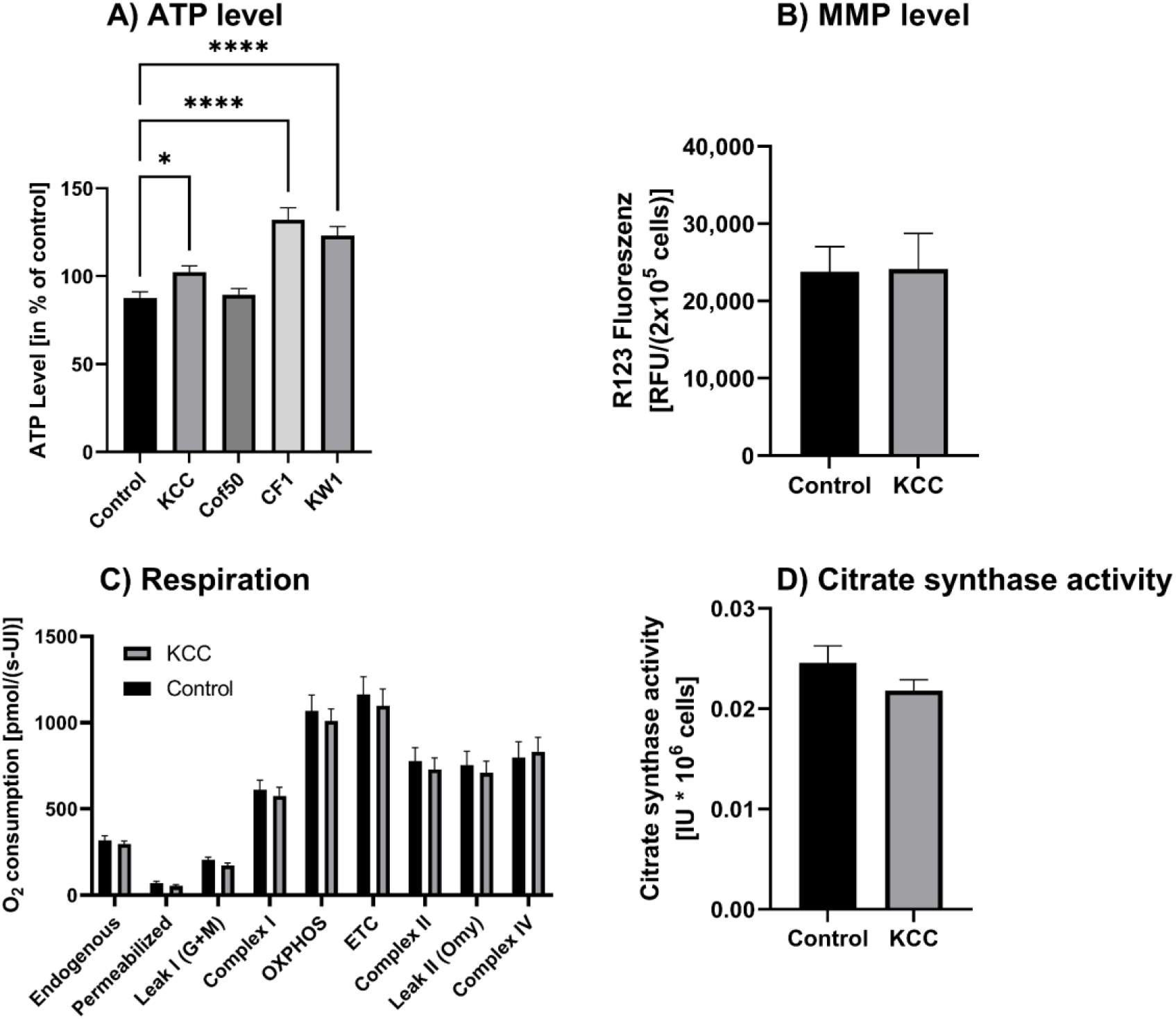
ATP level, MMP level, respiration and citrate synthase activity of SH-SY5Y- APP_695_ cells incubated for 24h with KCC. (A) ATP level of SH-SY5Y-APP_695_ cells incubated with KCC. Cells treated with cell culture medium served as control (100%). N = 8. B) MMP level of 2x10^5^ SH-SY5Y-APP_695_ cells incubated with N = 16. (C) Respiration of SH-SY5Y-APP_695_ cells incubated with KCC compared to the control.

**Figure 2:**
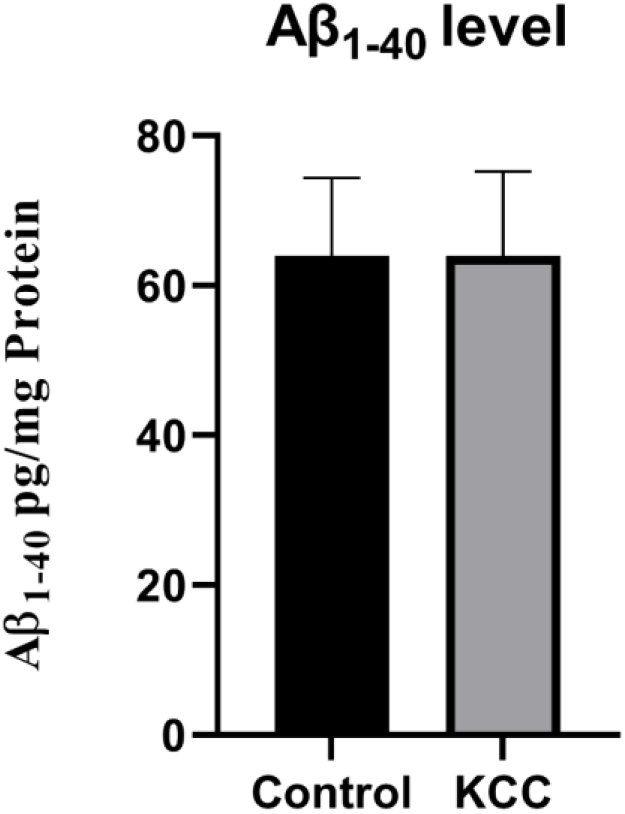
Effect of KCC on the Aβ_1-40_ level of SH-SY5Y-APP_695_ cells incubated for 24h. N = 10. Aβ_1-40_ levels were adjusted to the protein content. Significance was determined by Student’s unpaired t-test. Data are displayed as the mean ± SEM. KCC = kahweol 1µM, cafestol 1µM and caffeine 50µM.

Respiration under O2 consumption through the respiratory chain builds up the mitochondrial membrane potential, which is used by the ATP synthase to produce ATP. We measured the ATP level, MMP level, respiration and citrate synthase activity (Fig. 1). The KCC combination had a significant increasing effect (p = 0.0151) on the ATP level compared to the control (Fig. 1A). Also the single substances CF1 and KW1 (p = 0.0001) had a significant increasing effect compared to the control. Whereas the incubation with KCC had no effect on the MMP level, respiration or citrate synthase activity. Caffeine alone does not increase or decrease ATP levels. Only the combination with one or both diterpenes shows a significant increase at 50 µM (SupFigure 2). We chose the concentration of 50µM caffeine to be closer to the doses that are realistic when consuming coffee without damaging the cells or causing a loss of ATP.

SH-SY5Y-APP_695_ cells adjusted to international units (IU) of citrate synthase activity. N = 18. (G) Citrate synthase activity of SH-SY5Y-APP_695_ cells incubated with KCC compared to control. N = 18. Significance was determined by Student’s unpaired t-test. Data are displayed as the mean ± SEM. *P > 0.05, ****p> 0.0001. KCC = kahweol 1µM, cafestol 1µM and caffeine 50µM. Concentrations are given in µM

3.2 ***Aβ_1-40_ Production***

To measure the effect of KCC on the amyloid beta (Aβ) level SH-SY5Y-APP695 cells were incubated for 24h. Here we found no effect of KCC on the Aβ1-40 level compared to the control (Fig. 2).

### 3.3 Peroxidase

In order to investigated the effect of KCC or the single substances on the peroxidase activity in SH-SY5Y-APP695 cells we incubated them for 24h. Here we found no differences between incubations and the control with regard to the peroxidase activity (Fig. 3).

**Figure 3:**
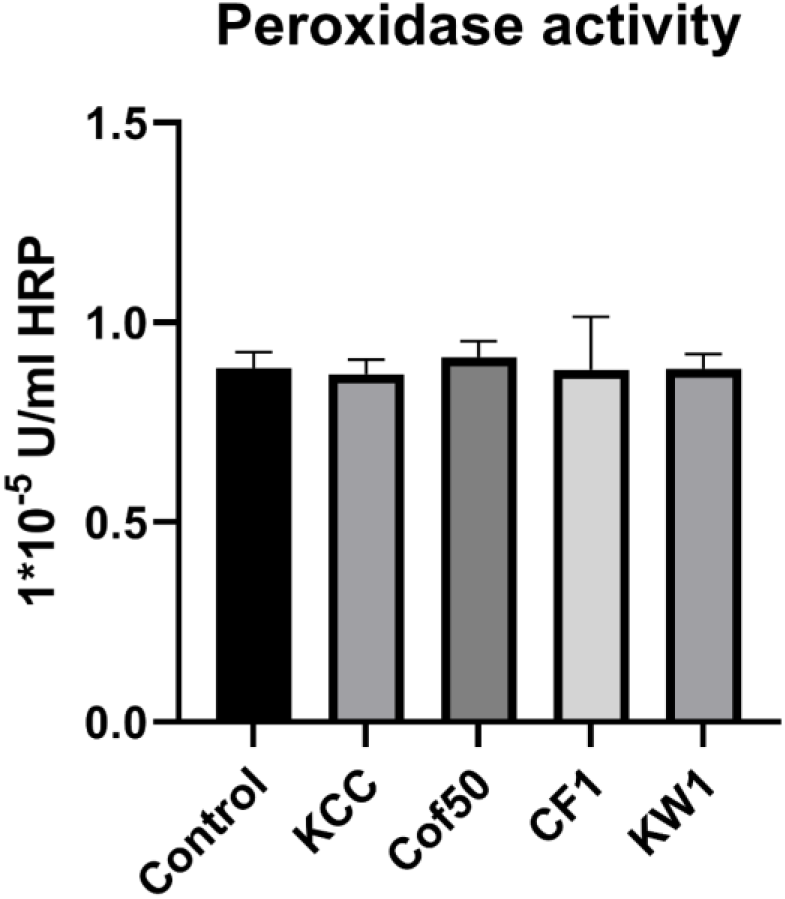
Effect of KCC and the single substances on the peroxidase activity of SH- SY5Y-APP_695_ cells incubated for 24h. N = 14. Significance was determined by Student’s unpaired t-test. Data are displayed as the mean ± SEM. KCC = kahweol 1µM, cafestol 1µM and caffeine 50µM. Concentrations are given in µM.

### 3.4 ROS measurement

In the next step, we examined the ROS level in SH-SY5Y-APP695 cells after the incubation with KCC or the single substances for 24h. Here, we found no different between both treatments (Fig. 4).

**Figure 4:**
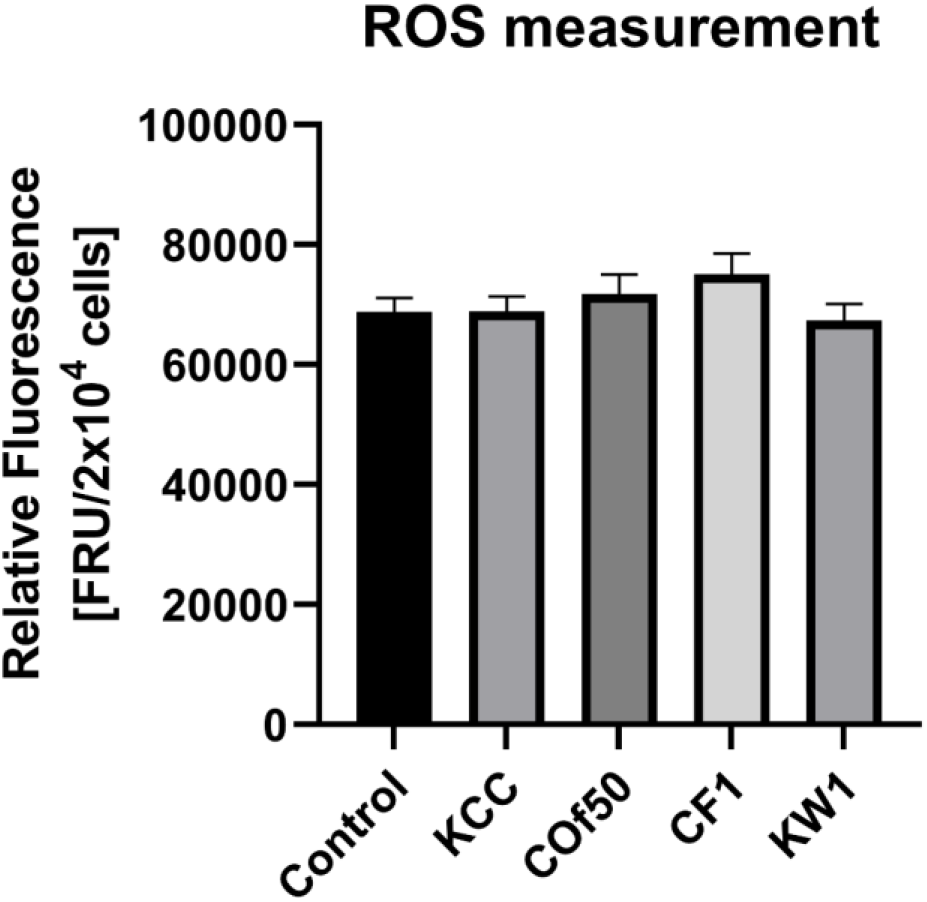
ROS measurement of SH-SY5Y-APP_695_ cells after incubation with KCC for 24h. N = 12. Significance was determined by Student’s unpaired t-test. Data are displayed as the mean ± SEM. KCC = kahweol 1µM, cafestol 1µM and caffeine 50µM. Concentrations are given in µM.

### 3.5 cAMP

To test the effect of KCC on the cAMP level, we incubated SH-SY5Y-APP_695_ cells for 24h. Here we had an increase, but not a significant, of cAMP level after the incubation with KCC compared to control (Fig. 5).

**Figure 5:**
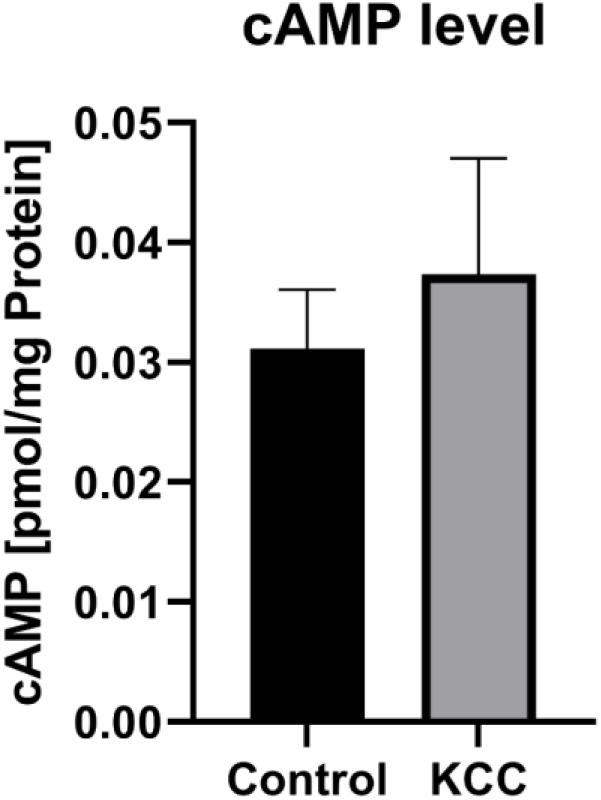
cAMP level of SH-SY5Y-APP_695_ cells after incubation with KCC for 24h compared to the control. N = 8. Significance was determined by Student’s unpaired t-test. Data are displayed as the mean ± SEM. KCC = kahweol 1µM, cafestol 1µM and caffeine 50µM.

### 3.6 Lactate and pyruvate level

To examine the effect of KCC on the lactate and pyruvate level we incubated SH- SY5Y-APP_695_ cells for 24h. There were no different between KCC compared to the control for the lactate level (Fig. 6A). However, the incubation with KCC significant increased the pyruvate value compared to the control (p = 0.0027) (Fig. 6B). Likewise, the lactate/pyruvate ratio (Fig. 6C) was significantly reduced by KCC compared to the control (p = 0.0141). To investigate whether the increased pyruvate levels were caused by a single substance. We determined the values of the individual substances. This showed that the pyruvate values were significantly increased by KW (p = 0.0062, Fig 6B). All the individual substances (Fig 6C) also significantly reduced the lactate/pyruvate ratio (Cof p = 0.0097, CF p = 0.0321, KW p = 0.0008).

**Figure 6:**
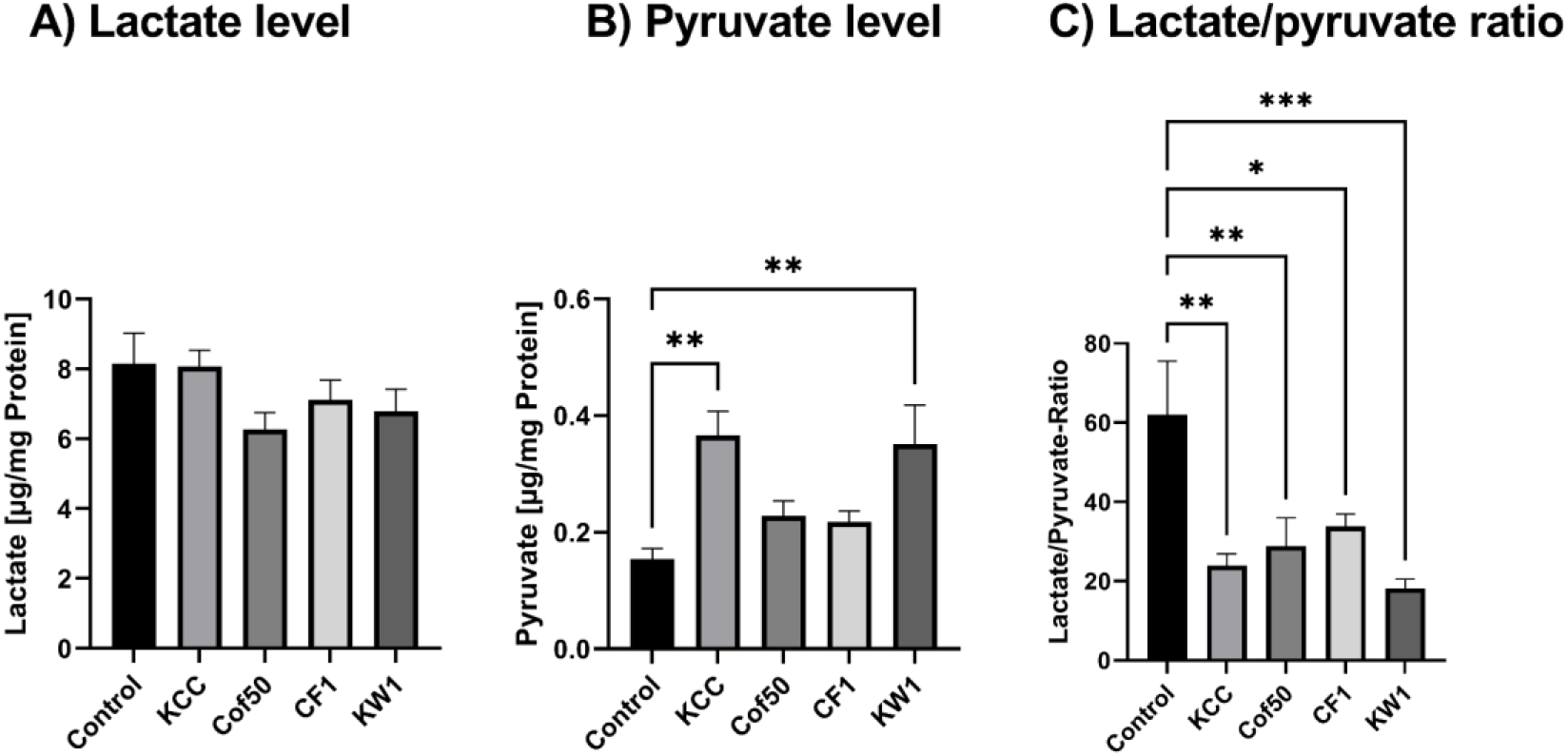
Effect of the on lactate and pyruvate level after 24h of incubation with caffeine (Cof), cafestol (CF) and kahweol (KW) or the combination KCC. (A) Lactate level of SH-SY5Y-APP_695_ cells after the incubation compared to the control. (B) Pyruvate level of SH-SY5Y-APP_695_ cells after the incubation with KCC compared to the control. (C) Lactate to pyruvate ratio. N = 8. Levels were adjusted to the protein content. Concentrations are given in µM. Significance was determined by one-way ANOVA. * p < 0.05 and ** p < 0.01, ***p > 0.001. Data are displayed as ±SEM. Concentrations are given in µM.

### 3.7 qPCR

To investigate the molecular basis of altered lactate and pyruvate levels, the gene expression of pyruvate dehydrogenase kinase 1 (PDK1) and lactate dehydrogenase A (LDHA) were examined after 24 h incubation using qRT-PCR. KCC had no significant effect on LDHA or PDK1 gene expression compared to control (Fig.7).

**Figure 7:**
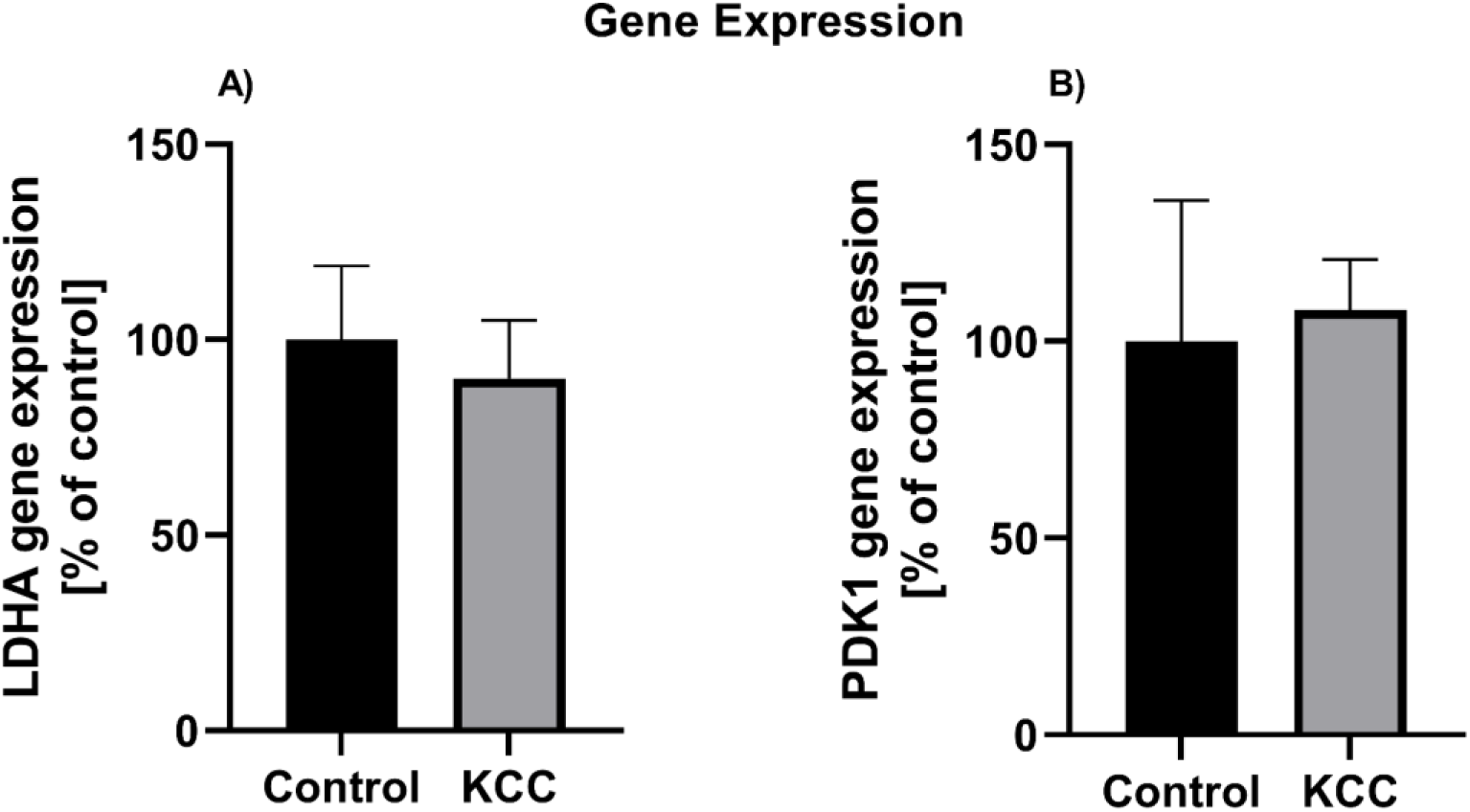
Effect on the incubation with KCC on the gene expression after 24h of incubation. (A) Gene expression of lactate dehydrogenase A (LDHA) of SH-SY5Y_695_ cells after the incubation with KCC compared to the control. (B) Gene expression of pyruvate dehydrogenase kinase 1 (PDK1) of SH-SY5Y_695_ cells after the incubation with KCC compared to the control. N = 8. The normalization factor was calculated based on the geometric mean of the levels of multiple control genes of *ß-actin (ACTB), glyceraldehyde-3-phosphate dehydrogenase (GAPDH)*, and *phosphoglycerate kinase 1 (PGK1)* according to the MIQE guidelines [33]. Significance was determined by Student’s unpaired t-test. Data are displayed as the mean ± SEM. KCC = kahweol 1µM, cafestol 1µM and caffeine 50µM.

## 4. Discussion

In the present work, we aimed to investigate different bioactive substances present in coffee. For this purpose, we selected three molecules that occur in larger concentrations in coffee - namely, caffeine, kahweol and cafestol. We tested various concentrations and combinations of these three compounds, of which 1µM kahweol, 1µM cafestol and 50µM caffeine (KCC) together were most promising. The compounds were tested in a SH-SY5Y-APP_695_ cells, an established model for early AD.

First, we investigated the effects of the single coffee compounds and its combination (KCC) on the mitochondrial dysfunction present occurring in SH-SY5Y- APP_695_ cells. The incubation with KCC significantly increased the ATP levels compared to the control (Fig. 1A). In addition, KW1 and CF1 increase the ATP levels significant. Caffeine alone does not increase or decrease ATP levels. Only the combination with one or both diterpenes shows a significant increase at Cof 50µM (SupFigure 2). In a transgenic mouse model of AD that oral treatment with caffeine intake of 0.6mg/d raised the ATP level significantly [34]. According to our result, kahweol was able to significantly rise ATP levels in SH-SY5Y cells a model [28]. It seems a combination of both compounds has equally positive effects on ATP levels. Due to the lack of knowledge it is unclear to what extent CF affects ATP levels. We found no indication that increased ATP levels were a result of enhanced MMP in consequence of an accelerated complex activity through the respiratory chain. In studies on SH-SY5Y cells with KW, no change in MMP or oxygen consumption could be measured by single administration, but KW was able to attenuate the effect by the stressor H_2_O_2_ [26,28]. This suggests that KW in combination with the other investigated substances has a positive effect on the maintenance of the MMP and the complex activity of the respiratory chain. In addition, the increased ATP level were not a result of a boosted mitochondrial mass (Fig. 1D), which leads to the assumption that these come from a different source. Another parameter related to mitochondrial quality and functionality are reactive oxygen species (ROS) that are produced in excess due to an insufficient proton transfer through the mitochondrial respiration chain. Due to their ability to interact with cellular components such mitochondrial membrane lipids or DNA, they can thus cumulatively cause oxidative damage that shortens lifespan and leads to impaired mitochondrial function [35]. However, in our study the tested compounds and their combination showed no antioxidant activity. Studies have shown that the addition of caffeine can lower ROS levels and thus have anti-inflammatory effects. Overall, caffeine is thought to reduce oxidative stress. Caffeine has been shown to inhibit the activity of NF-_k_B and increase that of superoxide dismutase [36]. CF is also thought to have a suppressive function of NF-_k_B [37]. Peroxidase, which catalyses the oxidation of organic and non-organic substances using H_2_O_2_, is considered an important antioxidant enzyme. In this process, it is supposed to protect against the harmful effects of physiological accumulation of ROS. Caffeine has been shown to increase the activity of antioxidant enzymes in the liver of mice or to increase overall antioxidant [38,39]. Glutathione peroxidase as well as superoxide dismutase, both antioxidant enzymes, benefit from increased activity after coffee addition in human experiments [40]. We examined the peroxidase activity that represents an important detoxification mechanism and found no significant differences by incubation with the single substances or its combination. The pathological appearance of AD is characterized by a deposition of oligomeric Aβ in the brain. For this purpose, we measured Aβ_1-40_ levels in our experiments (Fig. 2). There were no significant changes due to the incubations compared to the control. However, there is evidence that there is a caffeine-induced protective effect on the internalization of APP processing. Thus, Aβ_1-40_ concentration was significantly decreased due to the addition of caffeine [41]. In animal experiments with mice, it was shown that both a single and long-term administration of caffeine significantly reduced plasma Aβ levels. Increased cognitive performance and decreased Aβ deposition in the brain were also observed [42]. Studies in humans showed that moderate coffee consumption was associated with lower Aβ levels. This potential neuroprotective effect is consistent with other studies that suggest coffee consumers are at lower risk for AD [43–45]. It is possible that a longer incubation period with the three components would have had a greater effect on Aβ production.

Cyclic adenosine monophosphate (cAMP) belongs to the nucleotides and serves as a second messenger in the body. Its tasks are signal transmission in the context of metabolic pathways and hormonal effects, in which a signal cascade is induced by cAMP. Its main mode of action is to activate the cAMP-dependent protein kinase A enzyme family. One consequence of protein kinase A (PKA) being activated in this way is that it influences gene transcription by means of phosyphorylation of the cAMP response element binding protein (CREB) [14,15]. CREB affects neuron growth, neuron differentiation, synaptic plasticity and neurogenesis. In diseases such as Parkinson’s disease, Huntington’s disease and AD, an impairment of CREB has been observed [46]. The cAMP signaling cascades thus altered have the ability to modulate long-term memory. This is based on the interaction of phosphorylated CREB with target genes.

Thus, it is postulated that cAMP has a beneficial effect on cognitive performance in the context of short-term and long-term memory [47]. Cognitive performance deficits occurring in AD is mediated, among other things, by the cAMP/PKA/CREB signaling cascade. Aβ toxicity may represent one trigger of this impairment [17]. Furthermore, pathological Aβ concentration shows to cause a rapid and sustained decrease in PKA activity and inhibition of CREB phosphorylation [48]. Similarly, oxidative stress inhibits PKA activity and phosphorylation of CREB have. Administration of antioxidants has been shown to protect neurons from Aβ-induced inactivation by PKA [16]. In our study, Figure 5 shows an increase in cAMP levels by incubation with KCC compared with the control, although not a significant one. Studies have shown that caffeine has an increasing effect on cAMP concentration [49,50]. Furthermore, caffeine has shown a beneficial effect on PKA activity in animal models of AD [48]. Overall, many benefits lie in increasing the cAMP signalling cascade, which includes activation of CREB, sirtuin 1, and Nrf2. In the Nrf2 signalling pathway, potential modes of action of caffeine, cafestol, and kahweol overlap. Studies suggest that CF and KW may condition health-promoting effects via Nrf2, but unlike caffeine, without exerting influence on cAMP levels. There is no known cAMP modulation by CF or KW [51–53].

Glucose is the primary source of energy for brain activity. It is suspected that due to this, an impairment of glucose metabolism is associated with neurodegenerative diseases [54]. Lactate is the end product of anaerobic glycolysis catalysed by lactate dehydrogenase (LDH). Lactate is considered as an important bioenergetic metabolite formed in the absence of oxygen by fermentation or in the presence of oxygen by aerobic glycolysis [55]. Lactate is the link between the glycolytic and aerobic metabolic pathways [56]. It has been observed that when ATP production is impaired, lactate production for short- and long-term energy provision is stimulated. Thus, increased ATP levels may decrease cellular lactate demand [57]. In our work, despite increased ATP levels, there was no decrease in lactate values (Fig. 6A). Pyruvate is the end product of glycolysis and is considered a key molecule for numerous metabolic pathways. These include mitochondrial ATP generation and several biosynthetic pathways that cross the citrate cycle [58]. In general, neurons have an increased pyruvate demand due to a high energy turnover [59,60]. The significantly increased pyruvate concentration may indicates a relevant influence on the metabolism of pyruvate [61]. In our study, incubation with the combination of compounds significantly increased pyruvate levels (Fig. 6B). This effect might relay on KW, which as a single substance significantly increased the pyruvate levels, whereas the other single substances could only partly raise the levels. Other studies have shown that increased pyruvate levels can protect neurons from damage. In this case, the damage triggered by a stressor was annulated by an increased pyruvate concentration and at the same time pyruvate specifically protects the mitochondria from oxidative stress [62,63]. With regard to the lactate/pyruvate ratio (L/P), the results obtained indicate that there is a lowered ratio in favour of pyruvate. An increase in L/P is accompanied by an increase in NADH/NAD^+^. In our experiments, there should be a decreased ratio of NADH/NAD^+^ corresponding to L/P [56]. A shift in this ratio could have an impact on glycolysis, as there is inhibition of glycolysis when the NADH/NAD^+^ ratio is increased [56,64]. This may suggest that increased glycolysis may be present in the experiments performed here due to a decreased NADH/NAD^+^ ratio. Gene expression analysis of lactate dehydrogenase A (LDHA) and pyruvate dehydrogenase kinase 1 (PDK1) performed in our study (Fig. 7) showed no significant differences from control. There was a slight decrease in LDA and a slight increase in PDK1 expression. In an AD model with Drosophila melanogaster, downregulation of LDHA was shown to stimulate neuroprotective effects. Consistent with this, overexpression of LDHA showed a shortened lifespan as well as brain dysfunction. Based on this, LDHA can be considered a modulator of aging processes [65–67]. PDK1 inhibits pyruvate dehydrogenase (PDH) via reversible phosphorylation. PDH catalyses the oxidative decarboxylation of pyruvate to acetyl-CoA and CO_2_ in the mitochondrial matrix [63]. Shift in PDK activity may result in changes in mitochondrial function as well as neurological activity. With age, there is a decrease in PDH via increased PDK activity, which shifts energy production from aerobic to anaerobic glycolysis [68]. Overall, synergistically, the slight decrease in LDHA in combination with the slightly increased PDK1 could explain the increased pyruvate concentration. Thus, the increased ATP levels could be the result of increased glycolysis. This would go hand in hand with the observations made of the increased pyruvate concentration. Thus, an increased glucose metabolism up to the generation of pyruvate can be assumed. If an increased metabolism of glucose persists, need to be addressed by future studies.

## 5. Conclusion

In the present study, we observed the effect of a combination of different coffee components, kahweol, caffestol, and caffeine, on early AD symptomatology in a cell model of early AD. ATP levels were increase independent from mitochondrial function. Despite the lack of change in gene expression and the lack of change in mitochondrial complex activity, we suspect a change within glycolysis due to the change in pyruvate level. Further investigations have to address if coffee components affect glycolysis.

## Supplementary Materials

The following supporting information can be downloaded at:

## Author Contributions

investigation, L.B, M.T.L.; writing—original draft preparation, L.B.; supervision, G.P.E. All authors have read and agreed to the published version of the manuscript.

## Data Availability Statement

The data presented in this study are available on request from the corresponding author.

