## Supplement Data for "The Effect of Selected Coffee Constituents on Mitochon-Drial Dysfunction In An Early Alzheimer Disease Cell Model" (from another version of this preprint: version 1, 2023-04-20)

Supplemented Material for “THE EFFECT OF SELECTED COFFEE CONSTITUENTS ON EARLY ALZHEIMER SYMPTOLOGY IN A SH-SY5Y-APP_695_ CELL MODEL”

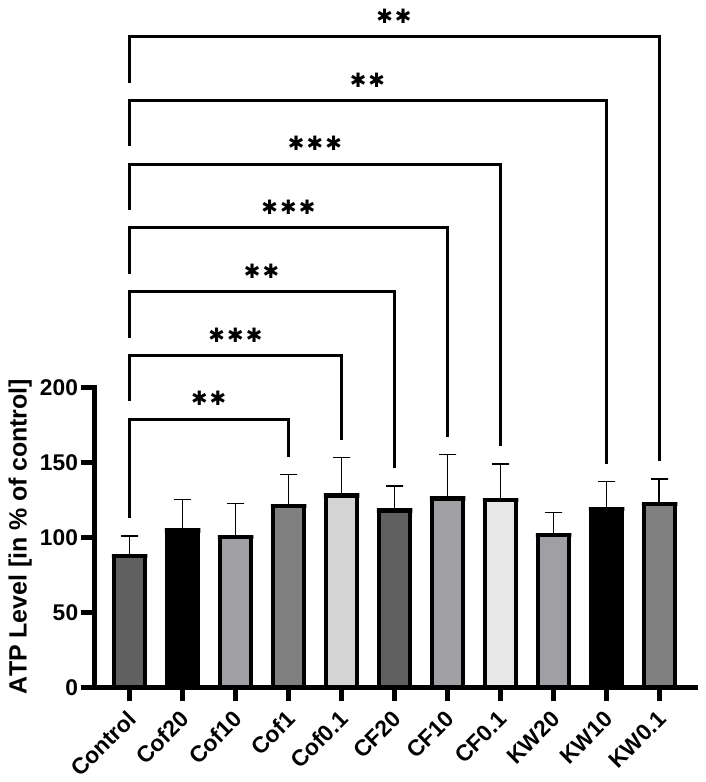

SupFigure 1: ATP level of SH-SY5Y-APP_695_ cells incubated for 24h with different concentration of caffeine (Cof), cafestol (CF) and kahweol (KW). Concentrations are given in µM. Cells treated with cell culture medium served as control (100%). N = 8. Data are displayed as the ± SD. Significance was determined by one-way ANOVA. **p > 0.01, ***p > 0.001, ****p > 0.0001.

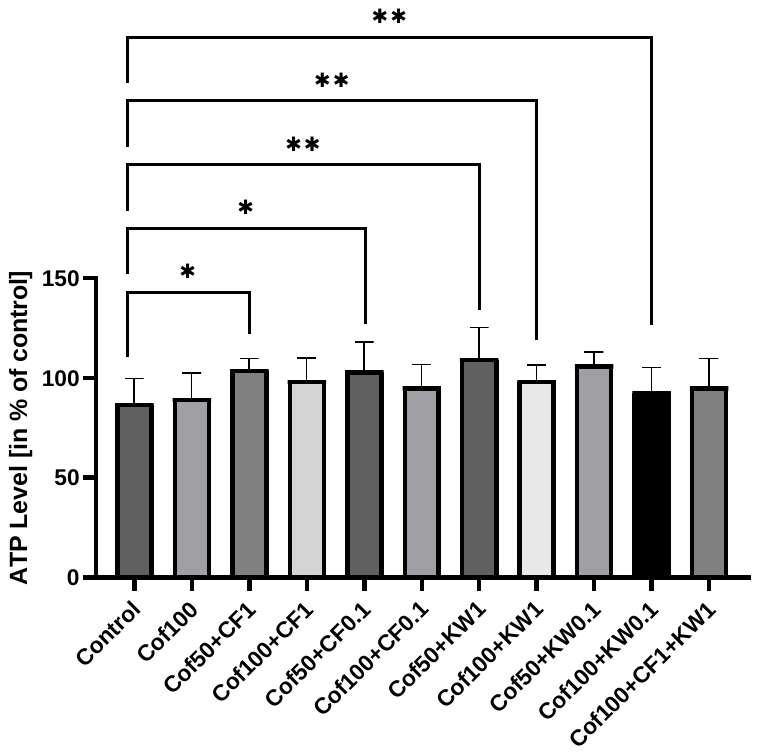

SupFigure 2: ATP level of SH-SY5Y-APP_695_ cells incubated for 24h with different concentration of caffeine (Cof), cafestol (CF) and kahweol (KW). Concentrations are given in µM. Cells treated with cell culture medium served as control (100%). N = 8. Data are displayed as the ± SD. Significance was determined by one-way ANOVA. *p 0.05, **p > 0.01.
